# STRIDE: accurately decomposing and integrating spatial transcriptomics using single-cell RNA sequencing

**DOI:** 10.1101/2021.09.08.459458

**Authors:** Dongqing Sun, Zhaoyang Liu, Taiwen Li, Qiu Wu, Chenfei Wang

## Abstract

The recent advances in spatial transcriptomics have brought unprecedented opportunities to understand the cellular heterogeneity in the spatial context. However, the current limitations of spatial technologies hamper the exploration of cellular localizations and interactions at single-cell level. Here, we present spatial transcriptomics deconvolution by topic modeling (STRIDE), a computational method to decompose cell types from spatial mixtures by leveraging topic profiles trained from single-cell transcriptomics. STRIDE accurately estimated the cell-type proportions and showed balanced specificity and sensitivity compared to existing methods. We demonstrate STRIDE’s utility by applying it to different spatial platforms and biological systems. Deconvolution by STRIDE not only mapped rare cell types to spatial locations but also improved the identification of spatial localized genes and domains. Moreover, topics discovered by STRIDE were associated with cell-type-specific functions, and could be further used to integrate successive sections and reconstruct the three-dimensional architecture of tissues. Taken together, STRIDE is a versatile and extensible tool for integrated analysis of spatial and single-cell transcriptomics and is publicly available at https://github.com/wanglabtongji/STRIDE.

## Introduction

The rapid development of high-throughput single-cell sequencing technologies(1-3) allows the investigation of cellular heterogeneity and the specificity of gene expression at an unprecedented resolution. However, recent reports indicate that cellular heterogeneity is not only modulated by the intracellular regulatory network, but also influenced by the extracellular microenvironment(4,5). Traditional bulk and single-cell RNA-sequencing (scRNA-seq) require the dissociation of tissues or the isolation of single cells, resulting in the loss of cell positions and their proximities to each other. The recent advances in spatial transcriptomics have enabled the measurement of gene expression while preserving the spatial information(6-10), which provide great opportunities to investigate the cellular heterogeneity(11), cell-cell communication(12), and the interplay between each other(13) in the spatial context.

Despite the existing applications of spatial transcriptomics on different tissues(12,14) and disease models(15,16), there remain many computational challenges. The cell-type assignment is one of the most important issues to be addressed, considering that other downstream analyses such as the detection of spatially restricted gene expression or cell-cell interactions highly rely on correct cell-type annotations. Many *in situ* capturing technologies, such as spatial transcriptomics (ST) and 10X Genomics Visium utilized spots with a diameter of 55∼100μm to record mRNA positions, and thus might cover multiple homogeneous or heterogeneous cells (1∼20 cells). Other high-resolution methods such as Slide-seq(9) and HDST(17) could improve the resolution to 1∼2 cells. However, the fixed beads or wells used in those techniques cannot match the cell boundaries perfectly, resulting in the overlap with several cells. As a consequence, the gene expression measured at one spatial location (i.e., spot or bead) can be regarded as a mixture of multiple cells. The within-spot heterogeneity greatly affects the performance of unsupervised clustering, which was usually used in the analysis of scRNA-seq.

To understand the cell type distribution from spatial transcriptomics, the most common strategy is to integrate it with scRNA-seq. There are two approaches for the integration of single-cell and spatial transcriptomics: gene signature scoring and deconvolution. Previous studies performed enrichment analysis using the gene signature derived from scRNA-seq(16,18), the utility of which is limited by the incomparability of the scores across different spatial locations or slides. On the contrary, the deconvolution methods aim to estimate the exact cell type proportions for each spatial location, either by applying regression models(9,19,20) or through fitting probability distributions(21,22). However, most of the existing deconvolution methods depend on the marker genes inferred from single-cell reference, which might suffer from the high drop-out rate and unwanted gene expression fluctuations. The topic model was initially proposed in the field of text mining to discover latent sematic structures from a large collection of documents. In recent years, it has been extended to bioinformatics to analyze single-cell epigenomics(23) and CRISPR screening data(24) owing to the tolerance to the sparsity of data and the interpretability of topics. Here we presented STRIDE, a topic-model-based deconvolution method for spatial transcriptomics by integrating with matched scRNA-seq. STRIDE first discovers cell-type-associated topics from annotated single-cell transcriptome by performing topic modeling. Then STRIDE applies the pre-trained topic model to infer the cell-type compositions for each location of spatial transcriptomics. By using simulated spatial transcriptomics data, we validated the power of STRIDE to predict the cell-type proportions in high accuracy and sensitivity. To demonstrate the broad utility of STRIDE, we applied it to three spatial transcriptomics datasets of different tissues including mouse cerebellum, human squamous cell carcinoma (hSCC), and human developing heart. We demonstrated that the topics generated by STRIDE could accurately reflect the spatial signatures of each cell type, improve the resolution of spatial clustering, and finally help in reconstructing the three-dimensional (3D) structures of tissue based on the integration of multiple slides.

## Materials and Methods

### Implementation of STRIDE

#### Discovery of topics from scRNA-seq by latent Dirichlet allocation (LDA)

Topic modeling is frequently used in the field of natural language processing to discover latent sematic structures, referred as topics, from a mass of collected documents. Latent Dirichlet allocation (LDA), a generative probabilistic model, is one of the most popular topic modeling methods(25). Here we utilize LDA for the discovery of functional topics from scRNA-seq data. In our model, given that there are *M* cells and *V* common genes the scRNA-seq data shares with the ST data, for the *m*th cell, the occurrences of topics are assumed to follow a multinomial distribution

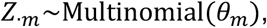

where *Z*_·*m*_ = (*Z*_1*m*_, *Z*_2*m*_, …, *Z*_*Km*_), represents the occurrences of all *K* topics in the *m*th cell, and *θ*_*m*_ is a *K*-dimensional parameter vector, which follows a Dirichlet distribution

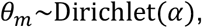

where *α* is a symmetric *K*-dimensional hyper-parameter, which denotes the prior weights of *K* topics in the *m*th cell. For the *k*th topic, the occurrences of genes are also multinomial-distributed

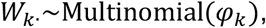

where *W*_*k*·_ = (*W*_*k*1_, *W*_*k*2_, …, *W*_*kV*_), represents the number of occurrences of all *V* genes in the *k*th topic, and *φ*_*k*_ is a *V*-dimensional parameter vector, which follows a Dirichlet distribution

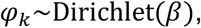

where *β* is a *V*-dimensional hyper-parameter, which denotes the prior weight of *V* genes in the *k*th topic. In STRIDE, an online variational Bayes (VB) algorithm(26) is applied to estimate the parameters and infer the gene-by-topic distribution and the topic-by-cell distribution, which is implemented by the python library Genism(27). With the output topic-by-cell distribution and user-provided cell-type labels, we can calculate the topic-by-cell-type distribution

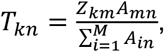

where *T*_*kn*_ represents the probability that the *n*th cell type has the *k*th topic, *Z*_*km*_ represents the probability that the *m*th cell has the *k*th topic, and *A*_*mn*_ = 1 *or* 0 means the *m*th cell belongs to the *n*th cell type or not. According to Bayes’ Theorem, the cell-type-by-topic distribution can be inferred as

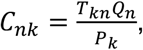

where *C*_*nk*_ represents the probability that the *k*th topic exists in the *n*th cell type, *Q*_*n*_ represents the probability of being the *n*th cell type (i.e., the percentage of the *n*th type of cells), and *P*_*k*_ represents the probability of the *k*th topic.

#### Selecting the optimal topic number

The number of topics makes a difference to the trained topic models. Theoretically, the identified topics should be representative and can distinguish different types of cells. To select the optimal topic number, STRIDE will run several models with different topic numbers and evaluate the models by the accuracy of single-cell re-annotation. To be specific, for each topic number, STRIDE will generate a cell-type-by-topic distribution *C*_*nk*_ as described above. Then, *SC*_*nm*_, the probability that the *m*th cell belongs to the *n*th cell type can be calculated as

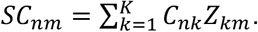

And each cell from scRNA-seq would be assigned the cell type with the largest probability. The cell-type prediction accuracy of single cells could be calculated by comparing the topic-derived cell type labels with the original labels users provide, and the topic model with the highest accuracy is selected to deconvolve ST data. If multiple models have the same highest accuracy, STRIDE will select the simplest one (i.e., the model with the smallest number of topics). The selection of the optimal topic number makes sure that the topic model STRIDE uses has the best ability to separate different cell types.

#### ST cell-type deconvolution

The gene expression measurements at a location in ST can be regarded as a mixture of multiple cells of different cell types. No matter what technology we use to characterize the cells, the latent topic structures (i.e., the gene-by-topic distribution) should be the same. With the gene-topic distribution derived from the first step, LDA could further estimate the topic distributions for each location in ST. That is,

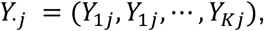

where *Y*_·*j*_ is the distribution of all *K* topics in the *j*th location. As in the single cell, the probability that the *j*th location belongs to the *n*th cell type can be calculated as

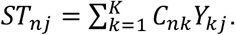

From another perspective, *ST*_*nj*_ could be viewed as the proportion of the *n*th type of cells at the *j*th location.

### Evaluating STRIDE’s performance using simulated ST data

#### Simulating ST data from BRCA scRNA-seq

To evaluate the deconvolution performance under different conditions, we generated three ST datasets from a breast cancer (BRCA) scRNA-seq dataset(28). To evaluate the accuracy of cell-type deconvolution, we simulated one ST dataset totally randomly. Each spot in the simulated ST dataset was a mixture of 2-10 cells randomly picked from scRNA-seq. The transcriptome profiles of selected cells were aggregated to represent the spot’s expression profile. To better mimic the real capture, if the synthetic spot had over 25,000 UMI counts, it would be down-sampled to 20,000 UMI counts. To demonstrate the robustness of STRIDE with different sequencing depths, we generated 1,000 spots with aggregated profiles of 10 randomly selected cells, and down-sampled them to 20,000, 15,000, 10,000, 5,000, 2,500 and 1,000 counts, respectively. Since the cell-type labels of selected cells are known, the resulting cell-type proportions can be regarded as the golden standard to evaluate the deconvolution performance.

In consideration of the co-localization of some cell types in reality, we generated an additional ST dataset from the same BRCA scRNA-seq data to simulate the spatially correlated cell-type distribution patterns in the tumor microenvironment. One typical tumor tissue could consist of three compartments: the tumor core (TC), the tumor stroma (TS), and the invasive margin (IM)(29). In addition to the three kinds of compartments, a special immune structure ---- tertiary lymphoid structure (TLS), which develops at the inflammatory sites in non-lymphoid tissues, was found to be present in several cancer types(30). Then we generated a ST dataset which simulated spots from TC, IM, TS and TLS, respectively. Spots from different regions have different cell-type co-localization patterns. For example, to simulate the TC region with infiltrating T cells, we generated 250 spots which contained primarily malignant cells and CD8 T cells. Detailed simulation rules are listed in Table 1. In addition, it was discovered that different infiltrating immune cells are limited to specific locations within or around a tumor (31). To mimic the true situation as much as possible, we also added some limitations of cell types included in different compartments (Table 1).

**Table 1.** Rules for simulation of the tumor microenvironment.

| Co-localization | Malignant ↔ CD8T | Malignant ↔ Fibro | Fibro ↔ CD4T/B | CD4T ↔ B |
| --- | --- | --- | --- | --- |
| Region<br>Cell type | Tumor core<br>(250 spots) | Invasive margin<br>(250 spots) | Tumor stroma<br>(125+125 spots) | TLS<br>(250 spots) |
| Malignant | 50% | 30% |  |  |
| CD4T | ✓ | ✓ | 30% | 30% |
| CD8T | 30% | ✓ | ✓ | ✓ |
| Tprolif |  | ✓ | ✓ |  |
| B |  | ✓ | Or 30% | 40% |
| Plasma |  | ✓ | ✓ | ✓ |
| Mono/Macro | ✓ | ✓ |  |  |
| Mast | ✓ | ✓ |  |  |
| Endothelial |  | ✓ | ✓ | ✓ |
| Fibroblasts |  | 30% | 50% |  |
| Myofibroblasts |  | ✓ | ✓ |  |

**Note:** ‘✓’ represents the existence of cell types with random cell-type fractions in different compartments, and the numbers represent the fixed percentages of correlated cell types. The correlation coefficient between correlated cell types was fixed as 0.7.

#### Function analysis on topics

LDA could discover latent topics from scRNA-seq, and each topic is composed of different genes with different probabilities. To ensure that the discovered topic profile would be able to characterize the cells, we performed functional enrichment analysis on the topics. For each topic, the top 200 genes with the highest probabilities were selected to perform gene ontology (GO) analysis by using clusterProfiler v3.18.1(32). For function analysis in mouse cerebellum dataset, biological process (BP) terms were form org.Mm.eg.db v3.12.0. In the human heart dataset, the top 50 genes for each topic were used to characterize the topics’ functions by enrichment analysis on the GO BP terms provided by org.Hs.eg.db v3.12.0.

#### Performance evaluation

As simulated data provided known cell-type compositions, we used two metrices, correlation and root mean squared error (RMSE), to assess the deconvolution performance. For each spot, Pearson’s correlation between the predicted cell-type proportions and the ground truth could be calculated to measure each spot’s deconvolution accuracy. We could also calculate Pearson’s correlation for each cell type to measure the performance in distinguishing different cell types. In addition, we used RMSE to evaluate the sensitivity and specificity. For each spot, according to the presence or absence, all the cell types were divided into two groups. RMSE was then calculated for each spot in the two groups separately. Specifically, in one spot, if RMSE is calculated only for cell types present in truth, it will measure the capability of each tool for identifying all true positives (i.e., sensitivity). On the contrary, if RMSE is calculated only for the absent cell types, it will measure how well they can identify the true negatives (i.e., specificity).

In the case when methods are applied to real ST data, there’s no golden standard to compare the prediction results with the truth. Instead, correlation between cell-type fractions and signature score was calculated to evaluate the performance. For each cell type, the top 50 specific marker genes, which were derived from *FindAllMarkers* function of Seurat v4.0.1, were defined as signature genes. The gene signature score was calculated for each spot, based on cell-type-specific signature genes through *AddModuleScore* function of Seurat.

#### Benchmarking different gene sets

To select the appropriate features for model training, we benchmarked the impact of different sets of genes. Since highly variable genes (HVGs) are useful for cell clustering in scRNA-seq analysis, we took HVGs into account. In general, all genes, marker genes, HVGs, and combination of marker genes and HVGs were used as the input for STRIDE separately. For each gene set, the optimal number of topics was selected according to the accuracy of cell-type assignment for single cells. Spot-level Pearson’s correlation was utilized to evaluate the performance of models with different gene sets.

#### Benchmarking different deconvolution methods

We used the simulated ST data to compare the performance of STRIDE with other ST deconvolution tools, including SPOTlight v0.1.6, NMFreg, Seurat v4.0.1 CCA, RCTD v1.2.0 and Cell2location v0.5. Each published method was run with all the parameters set as default values following their documentations. Methods were evaluated using correlation and RMSE as described in ‘Performance evaluation’ section.

### Application of STRIDE on Slide-seq V2 mouse cerebellum data

#### Collection of mouse cerebellum single-cell and spatial transcriptomics data

Due to the well-studied structure and complex cell-type composition, mouse cerebellum can be used as a model tissue to assess the cell-type deconvolution performance naturally. Slide-seq V2 mouse cerebellum data was obtained from a previous study(21). The single-nucleus mouse cerebellum data collected by snRNA-seq protocol was derived from another transcriptomic atlas study(33), which provided detailed cell-type annotation information.

#### Cell-type deconvolution of mouse cerebellum data

The single-cell and spatial gene count matrices were scaled by the UMI counts in each cell or spot to avoid the impact of different cell types. Then we performed unit-variance normalization by gene to standardize the expression levels of different genes. The top 500 differentially expressed (DE) genes of each cell type were selected as feature genes to run STRIDE. According to the prediction accuracy of single cells in scRNA-seq, the model with 40 topics was selected to perform deconvolution on spatial data. Given that the resolution of Slide-seq V2 is up to 1∼ 2 cells, each pixel in spatial data was assigned the cell type with the largest probability. To reveal the anatomical features clearly, the granule cell region was cropped out from the whole slide as the original study did. And only the 7 of 19 most common cell types were displayed in Figure 3.

#### Topic and marker visualization

STRIDE could discover latent topics and estimate the topic profile of spatial locations through LDA. The topics were proved to be associated with specific cell types and have cell-type-related functions by GO analysis. To visualize the spatial distribution of topics, we selected four cell types, oligodendrocytes, granule cells, Purkinje cells, and molecular layer interneurons. For each cell type, the probability distribution of representative topics was displayed. If a cell type had multiple associated topics, the sum of these topics was used to represent the cell type. For comparison, the distribution of marker genes was visualized. The top 100 DE genes of each cell type were defined as marker genes. The sum of the expression of marker genes was calculated and used as the cell type’s signature score.

### Application of STRIDE on human squamous cell carcinoma ST dataset

#### Collection and preprocessing of SCC scRNA-seq and ST data

The human SCC ST dataset was obtained from a published study on tumor microenvironment. The study also provided matched scRNA-seq data of the 10X Genomics Chromium platform. In our study, we only used the sample, replicate 2 of patient 2. The scRNA-seq data of patient 2 was clustered and annotated using the markers provided by the original publication. As one patient only had limited number of cells, the myeloid cells were classified into two clusters, DC and Mono/Macro (monocytes and macrophages).

#### Cell-type deconvolution of human SCC ST data

To run STRIDE, the single-cell and spatial gene count matrices were scaled and normalized in the way described in the mouse cerebellum dataset. The top 500 marker genes were identified for each cell type in scRNA-seq, and were used to train the topic models. The model with 28 topics was selected automatically to be the optimal one. The cell-type deconvolution result of ST was visualized by the scatter pie plot.

#### Deconvolution-based spatial clustering

The cell-type deconvolution could provide high-resolution view of cell-type composition. Spots with similar cell-type composition are assumed to belong to the same cluster. By combining cell-type composition similarity with the additional spatial information provided by ST, spatial domains could be further identified. To identify the spatial domains, k-means clustering was performed based on both the spot’s own cell-type composition and its surrounding cell populations. In the case of SCC, equal weights were given to the two factors, and k was set to 6 to run k-means clustering. C3 cluster was defined as tumor-core region, and spots surrounding the C3 cluster were defined as the tumor-edge region, which mainly consisted of C1, C4 and C5 spots. To divide the region outside tumor into the stroma and the tumor-stroma border, for each spot except C3, we calculated the distance between the spot and its nearest C3 spot, and regarded it as the distance to the tumor region. Then, spots with distance longer than 4 and shorter than 4 were defined to be the stroma region and tumor-stroma border, respectively.

#### Deriving high-resolution gene expressions

STRIDE could estimate the topic profiles for both scRNA-seq and spatial transcriptomics. Then the correspondence between single cells in scRNA-seq and spots in ST could be constructed according to the similarity of topic distributions. Here we calculated the cosine similarity to measure the similarity between spots and cells. For each spot in ST, according to the spot-cell similarity, 10 most similar cells from scRNA-seq were selected in proportion to cell-type composition inferred by STRIDE. In this way, the gene expression in each spot can be dissected into cell-type-specific expression.

#### Function analysis of epithelial cells in different clusters

The spot-matched single cells selected from the last steps could be pooled together to study the cellular heterogeneity with high resolution. To compare the difference of epithelial cells in different regions, we extracted epithelial cells predicted to be located in tumor-core and tumor-edge for differential analysis. In order to characterize the functions of region-specific epithelial cells, we performed enrichment analysis on hallmark pathways and GO BP terms collected from the Molecular Signatures Database (MSigDB v7.1)(34) and org.Hs.eg.db v3.12.0, respectively.

### Application of STRIDE on developing human heart ST dataset

#### Collection and preprocessing of human heart scRNA-seq and ST data

The developing human heart data were derived from a previous study(14). The study provided 4, 9 and 6 heart sections from three timepoints, 4.5-6, 6.5, and 9 PCW, respectively, which were profiled by ST technology. They also provided a 6.5 PCW cell-type map with single-cell resolution constructed by smFISH. The single-cell transcriptional profiles of the 6.5 PCW tissue sample were generated through 10X Genomics Chromium platform. The scRNA-seq data was clustered and annotated using the markers provided by the original study. Notably, three fibroblast-like cell clusters were combined into a large fibroblast-like cluster in our study.

#### Cell-type deconvolution of human heart ST data

To run STRIDE on the human heart dataset, the top 100 DE genes of each cluster compared to all other cells were identified by Wilcoxon test. Because epicardium-derived cells, smooth muscle cells, and fibroblast-like cells are transcriptionally similar, the marker genes of each of the three cell types were identified by comparing one with the other two. Markers genes for atrial and ventricular cardiomyocytes were discovered in the same way. The scaled and normalized count matrices with marker genes were utilized to train the topic models. The model with 28 topics was further employed to deconvolve the ST data. The deconvolution results of sample 3, 9 and 16 were visualized by the scatter pie plot.

### 3D model construction of human heart

#### Slide alignment based on STRIDE-derived topics and spatial structure

The human heart samples for each timepoint were sectioned along the dorsal-ventral axis, which can be used to reconstruct the spatial architecture of developing human hearts. Here we modified a published method PASTE(35), which utilized Fused Gromov-Wasserstein Optimal Transport (FGW-OT) algorithm, to integrate multiple tissue layers from a ST experiment. The original PASTE aligns and integrates adjacent layers based on the transcriptional and spatial similarity. Here we made a little change to the goal problem that we expected to find a mapping between spots from two layers which had similar topic profiles and preserved spatial relationship. The problem was formulated by finding a mapping 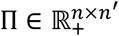 that minimized the following transport cost function:

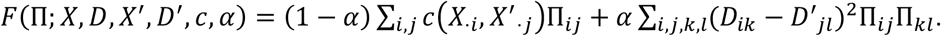

(*X, D*) and (*X*′, *D*′) denote two layers of one ST experiment, where 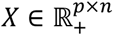 and 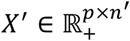 represent topic profiles of the *p* topics in *n* and *n*′ spots, and 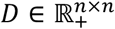 and 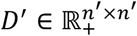 represent the spot distance matrices within each layer. The parameter *α* ∈ [0, 1] is used to weigh up the topic distribution similarity and spatial distance similarity. If *α* = 0, only topic distribution similarity is taken into consideration, and vice versa. The problem was solved by Python Optimal Transport (POT) library.

#### 3D reconstruction of PCW6.5 human heart

To reconstruct the 3D structure of the 6.5 PCW human heart, we sequentially aligned multiple adjacent ST layers. For a series of sequential layers (*X*^(1)^, *D*^(1)^), …, (*X*^(9)^, *D*^(9)^) from the dorsal to the ventral, we found the mapping Π^(*k*)^ (*k* = 1, …, 8) between the *k*th and (*k* + 1)th layer. Based on the mappings, we should project all tissue layers to the same spatial coordinate system. This is a generalized weighted Procrustes problem. That is, to find a translation vector 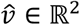 and a rotation matrix 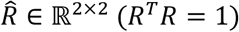 that minimize the weighted distances between matched spots

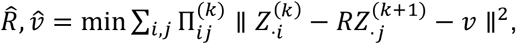

where *Z*^(*k*)^ ∈ ℝ^2×*n*^ represents the spot coordinate matrix of the *k*th layer. The problem was solved by using SVD (see PASTE(35) for more details). For Figure 6A, only three tissue sections, sample 8, 9 and 10 were displayed to visualize the alignment and integration result. For Figure 6B, all nine ST slides from 6.5PCW were displayed to provide a general view of the reconstructed 3D structure of the human heart.

## Results

### STRIDE: a topic-model-based method to deconvolve spatial transcriptomics using scRNA-seq

STRIDE is designed to deconvolve the cell-type composition of spatial transcriptomics locations by integrating scRNA-seq data from the same tissue (Figure 1). We assumed that transcriptional measurements obtained from spatial transcriptomics and scRNA-seq of the same tissue have common cell types with similar features, and thus can be projected to a common latent space. Here, we characterized the common latent space by topic modeling, a statistical model originally used in natural language processing. The first step of our method is to infer the cell-type-associated topic profiles from scRNA-seq. Topic modeling on cell-by-gene matrix from scRNA-seq data could generate two probability distributions. One is the probability of a gene belonging to a topic (i.e., gene-by-topic distribution). In addition, topic modeling also estimates the contributions of different topics for each cell (i.e., topic-by-cell distribution), which could be summarized to the cell-type-by-topic distribution using Bayesian method. Next, the pre-trained topic model could be employed to infer the topic distributions of each spot in spatial transcriptomics. Finally, STRIDE could estimate the cell-type fractions of each spot by combining cell-type-by-topic distribution and topic-by-spot distribution.

**Figure 1.**
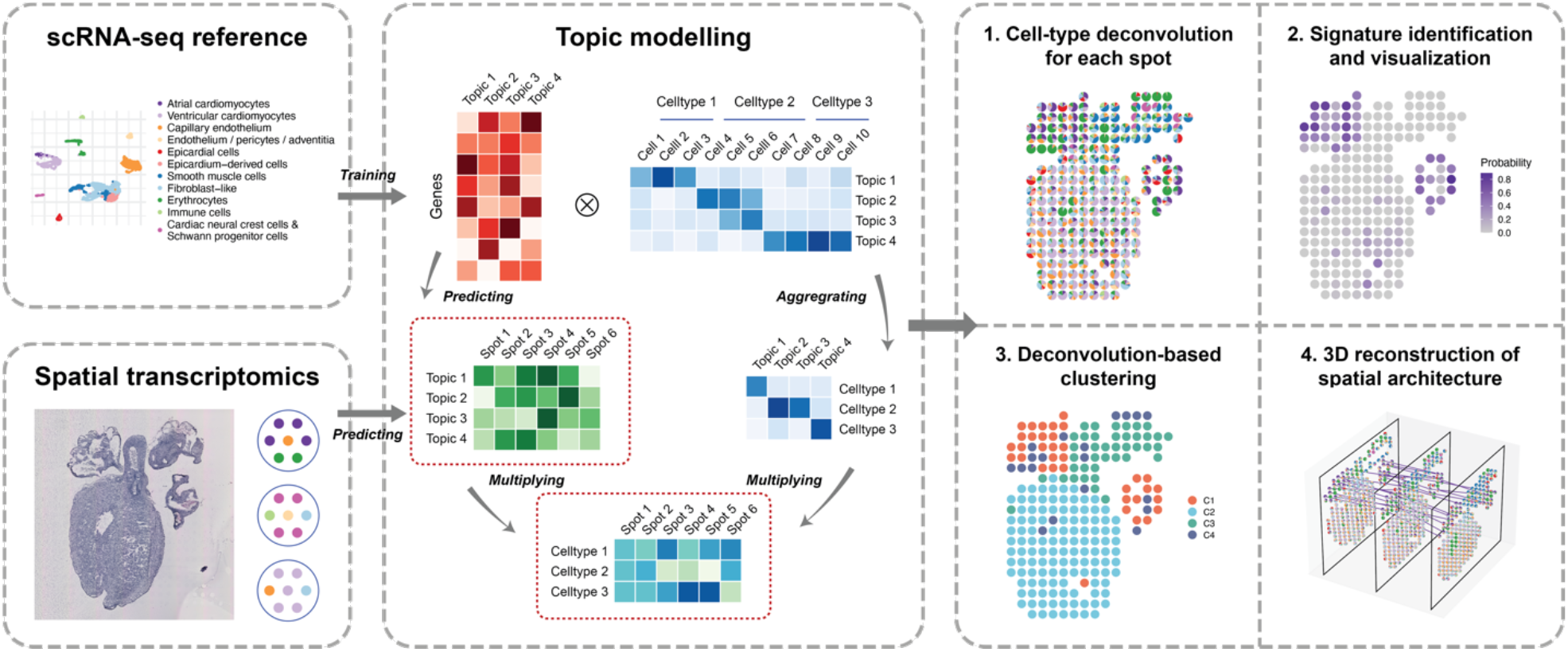
Schematic representation of the STRIDE workflow. First, STRIDE estimates the gene-by-topic distribution and the topic-by-cell distribution from scRNA-seq. The topic-by-cell distribution is then summarized to the cell-type-by-topic distribution by Bayes’ Theorem. Next, the pre-trained topic model is applied to infer the topic distributions of each location in spatial transcriptomics. By combining cell-type-by-topic distribution and topic-by-location distribution, the cell-type fractions of each spatial location could be inferred. STRIDE also provides several downstream analysis functions, including signature detection and visualization, spatial domain identification and reconstruction of spatial architecture from sequential ST slides of the same tissue.

Besides the cell-type composition deconvolution for each spot, STRIDE also provides several downstream analysis functions, including (1) signature (i.e., topic) detection and visualization, (2) spatial clustering and domain identification based on neighborhood cell populations and (3) reconstruction of three-dimensional architecture from sequential ST slides of the same tissue. The methodology details are described in the method section and the examples are shown in the following results.

### STRIDE yielded more accurate cell-type composition estimates on simulated ST data than existing tools

To evaluate the performance of STRIDE, we generated synthetic mixtures of cells with known cell-type labels. Specifically, we simulated three spatial transcriptomics datasets from a breast cancer (BRCA) scRNA-seq dataset(28) to imitate different conditions. For each simulated location, single cells from scRNA were selected randomly or by fixed cell type proportions and their transcriptomic profiles were combined to synthesize a mixture. The synthetic mixtures with known cell-type compositions could serve as ground truth to benchmark the performance of STRIDE on decomposing cell types.

We first validated the ability of topic modeling to discover cell-type-specific topics. We derived 28 different topics which were enriched in distinct cell types (Figure 2A), indicating the association between topics and specific cell types. We also performed Gene ontology (GO) function analysis on top enriched genes for each topic (Supplementary Figure S1A). GO biological process (BP) terms related to tumor metastasis, such as platelet aggregation, were detected in malignant-cell-associated topic (Topic 22), whereas immune-related functions were enriched in immune-cell specific topics (Supplementary Figure S1B). Furthermore, the B-cell, T-cell and myeloid cell enriched topics had distinct enriched terms, suggesting that the topics generated by STRIDE could accurately separate different immune cell types (Supplementary Figure S1B). In addition, when we validated the trained topic model using the same scRNA-seq dataset used for training, STRIDE achieved high cell-type assignment accuracy (87.13%, n = 33043 cells) (Figure 2B, see Materials and methods). Cell types such as CD4^+^ and CD8^+^ T cells, which are similar in their transcriptome, accounted for most of the errors (79.51%, n = 4251 errors).

**Figure 2.**
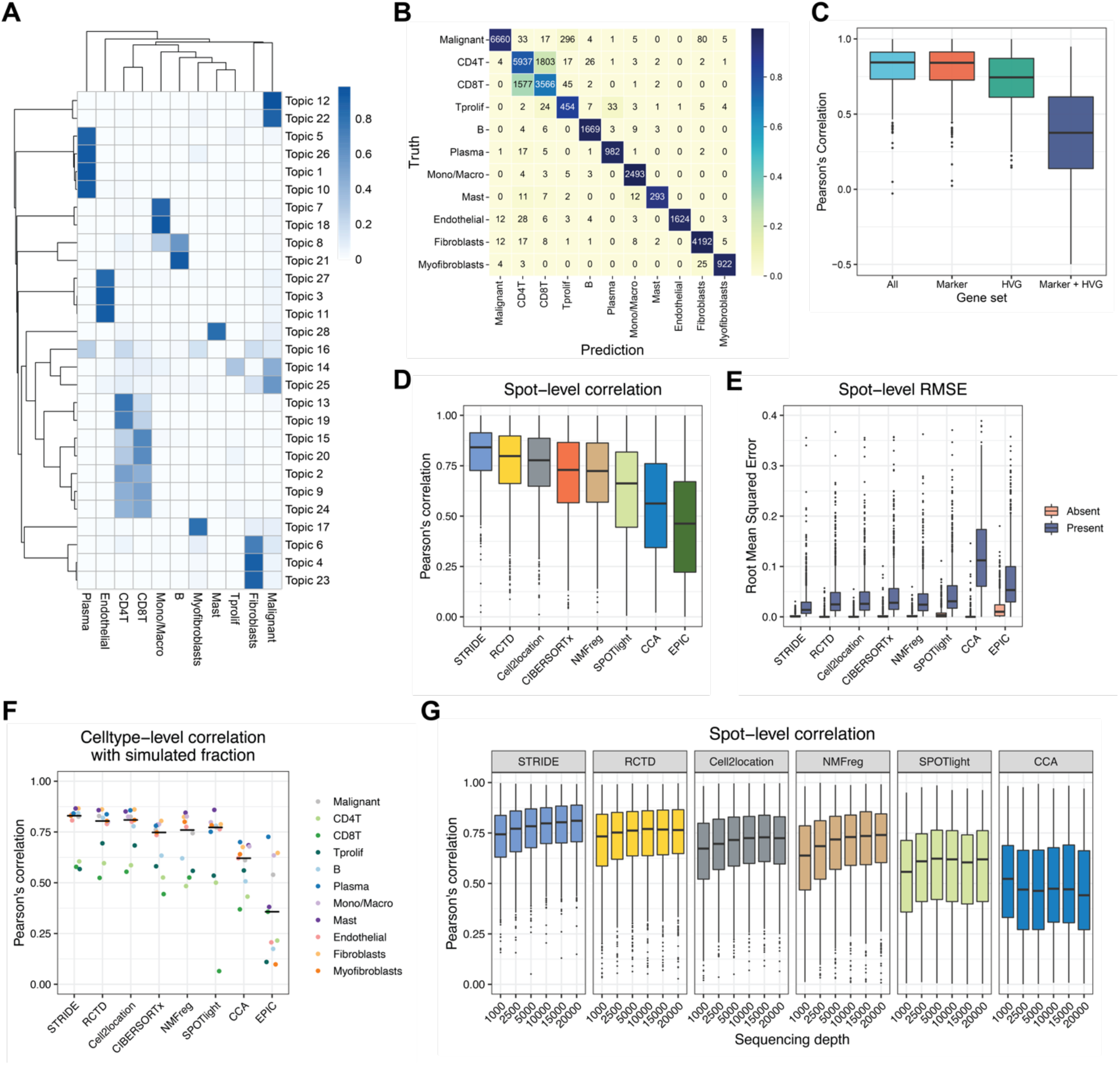
Benchmarking STRIDE’s performance using simulated data. A. The cell-type-by-topic distribution estimated by STRIDE. The color represents the probability that one topic exists in one given cell type. B. Validation of the trained topic model on the scRNA-seq used for training. The confusion matrix reflects the consistency between the prediction and the truth for the cell-type assignment of scRNA-seq data. The value represents the number of cells that belong to one cell type and are predicted to all different cell types. The color represents the proportion of cells belonging to the cell type on the y axis and classified as the cell type on the x axis. C. Benchmark of STRIDE’s performance on different gene sets. The box plot reflects the distribution of Pearson’s correlation calculated between the predicted cell-type proportion and the ground truth for each spot. D. Benchmark of STRIDE’s accuracy against different deconvolution methods. The box plot reflects the overall distribution of Pearson’s correlation calculated in each spot for each method. E. Benchmark of STRIDE’s sensitivity and specificity against different deconvolution methods. In each simulated location, the cell types were divided into two groups according to the presence (blue) and absence (pink), and RMSE was calculated within each group separately. The box plot reflects the distribution of RMSE in different methods. F. Benchmark of the ability to distinguish diverse cell types across different deconvolution methods. Pearson’s correlation between the predicted proportions and the ground truth was calculated for cell type. The black line in each column indicates the median of different cell types’ correlation for each method. G. Benchmark of STRIDE’s robustness against different deconvolution methods on the simulated dataset with different sequencing depths.

Feature selection is usually an important step to achieve a balance between resource consumption and model performance in machine learning algorithms. We then benchmarked the impact of feature gene selection on STRIDE’s deconvolution performance using simulated data. We calculated the overall Pearson’s correlation coefficients between estimated cell-type proportions and real proportions within each spot for evaluating the performance. Interestingly, model trained using marker genes (defined as DE genes for each cell type) showed comparably decent performance with model trained with all genes, while the models trained using highly variable genes (HVGs) or markers plus HVGs showed poor performance (Figure 2C, Supplementary Figure S1C). Therefore, STRIDE selects marker genes for model training by default, but also allows users to provide more specific cell-type signatures.

Next, we compared the performance of STRIDE with other published cell-type deconvolution tools, including methods developed for spatial transcriptomics, such as SPOTlight(19), NMFreg(9), Seurat CCA(36), RCTD(21), and cell2location(22), as well as the ones for bulk RNA-seq, such as CIBERSORTx(37) and EPIC(38). STRIDE showed the highest concordance between the prediction and the ground truth, and RCTD and cell2location showed slightly worse consistency (Figure 2D). To further evaluate the sensitivity and specificity, in each spatial location, we divided all cell types into two groups according to the presence and absence, and calculated root mean squared error (RMSE) within each group (see Materials and methods). STRIDE achieved a balance between sensitivity and specificity, while other methods such as CCA, RCTD and cell2location, achieved high specificity at the expense of low sensitivity (Figure 2E). We also assessed the methods’ capability to distinguish different cell types. STRIDE exhibited the overall best performance among all methods (Figure 2F, Supplementary Figure S1D). All of these methods showed poor performance in discriminating three T cell subtypes (i.e., CD4^+^ T cells, CD8^+^ T cells and proliferating T cells), which shows highly similar expression profiles even in the scRNA-seq data. Considering that in reality some cell types are co-localized, we generated an additional ST dataset from the same BRCA scRNA-seq data to simulate the spatially correlated cell-type distribution patterns in the tumor microenvironment. Among all the methods, STRIDE displayed generally best performance, in terms of both the concordance with the ground truth, and the balance between sensitivity and specificity (Supplementary Figure S1E-G). Lastly, in the real scenario some spots might be partially captured, and the sequencing depth might affect the number of captured genes as well as the deconvolution results. We further benchmarked the performance of STRIDE with different sequencing depths. Although the deconvolution performance declined with the decrease of the sequencing depth, STRIDE was still more robust to the sequencing depth compared to other methods (Figure 2G). Collectively, these results demonstrated that STRIDE could estimate the proportions of diverse cell types with high accuracy, and are tolerated to spatially co-localized cell-type distributions as well as low sequencing depth.

### Application of STRIDE on mouse cerebellum dataset revealed spatial localization patterns of cell types and novel spatial signatures

To demonstrate STRIDE’s usability on real spatial transcriptomics data, we applied it on a mouse cerebellum section profiled by Slide-seq2(21). The mouse cerebellum presents clearly defined cell-type layer structures, and thus can be used to assess the cell-type decomposition performance. We collected a published snRNA-seq dataset as reference(33), and applied STRIDE to map reference cell types to spatial locations of Slide-seq2 image (Figure 3A). Consistent to the prior literature (Figure 3B)(39), the two types of molecular layer interneurons, MLI1 and MLI2, were mapped to the top and outermost layer of the cerebellar cortex (i.e., molecular layer). Bergmann and Purkinje cells were colocalized to the same middle layer, Purkinje layer, while granule cells were localized to the bottom layer, granular layer. The oligodendrocytes and astrocytes were scattered below the granular layer (Figure 3A). Taken together, STRIDE could accurately deconvolve the cell types and reconstruct the layered structure of the mouse cerebellum.

**Figure 3.**
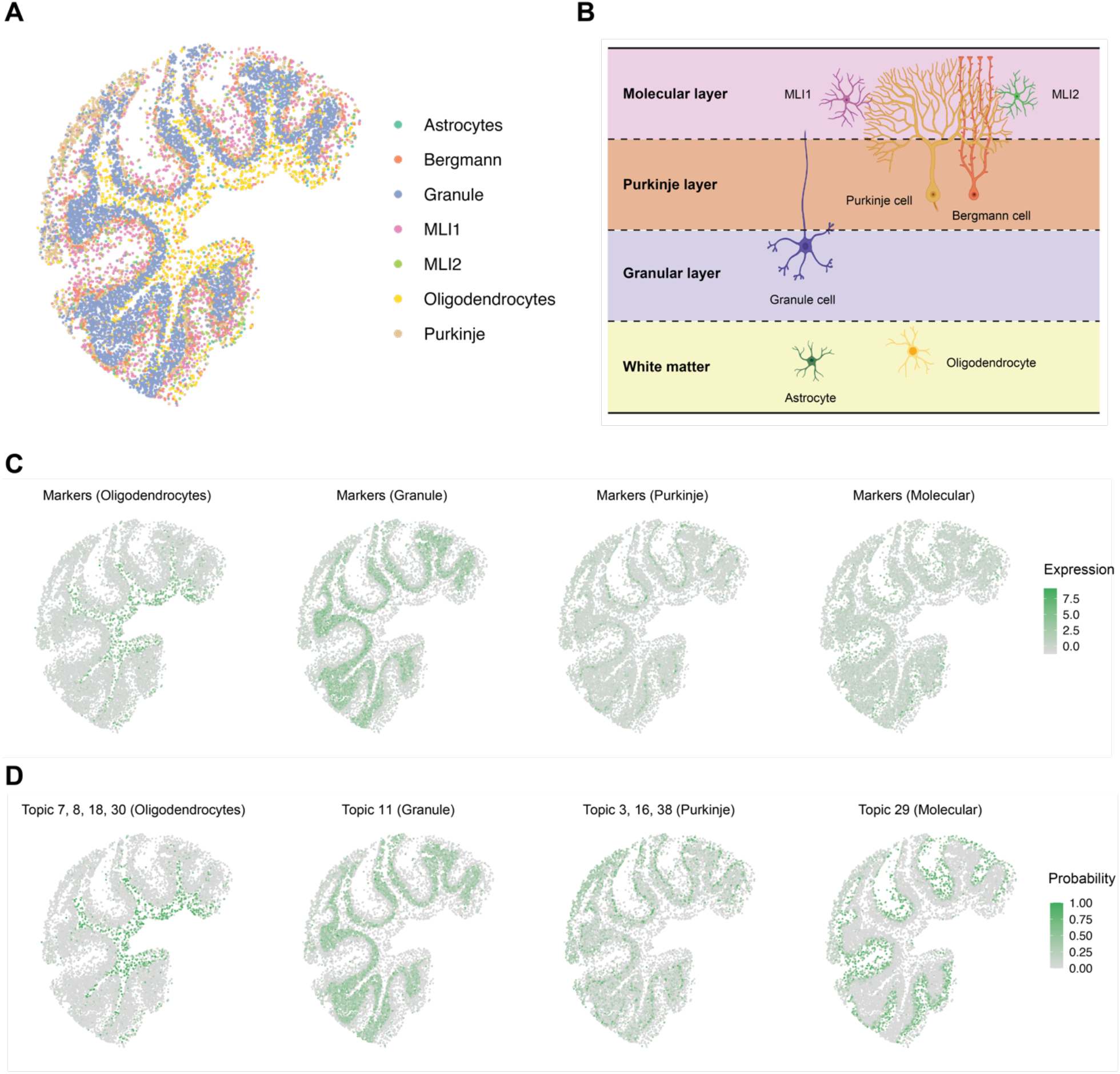
Application of STRIDE on the mouse cerebellum. A. The spatial distribution of the 7 most common cell types predicted by STRIDE. Each point represents the pixel captured by Slide-seq V2, and colors represent different cell types. B. The schematic of the layered structure of the mouse cerebellum. The cerebellum cortex is divided into three layers. At the top lies the molecular layer which contains two types of molecular layer interneurons, MLI1 and MLI2, along with the dendritic trees of Purkinje cells. At the middle lies the Purkinje layer which contains the body of Purkinje cells and Bergmann cells. At the bottom lies the granular layer which contains the granule cells. Below the cortex is the region of white matter enriched with oligodendrocytes and astrocytes. (Created with BioRender.com.) C. The expression patterns of cell-type-specific marker genes for oligodendrocytes, granule cells, Purkinje cells and MLIs. The color represents the summed expression values of top 100 marker genes. D. The distribution of cell-type-associated topics for oligodendrocytes, granule cells, Purkinje cells and MLIs. The color represents the summed probability of associated topics in each pixel.

Although there’s no ground truth for real spatial transcriptomics data, we could potentially compare the deconvolution result achieved from STRIDE with the cell-type-specific markers. We found that the spatial distribution of cell types inferred by STRIDE was generally consistent with the spatial distribution of marker genes (Figure 3C). For example, the marker genes of oligodendrocytes were highly expressed in the white matter underneath the gray matter of the cerebellar cortex, where oligodendrocytes are highly enriched (Figure 3C). In line with expectations, the markers for granule cells, Purkinje cells and molecular layer interneurons were distributed from the inner layer to the middle layer, and thence to the outermost layer, respectively (Figure 3C). The further correlation analysis between gene signature score and cell-type proportion revealed roughly corresponding relationships, despite the confusion between astrocytes and Bergmann cells, and between MLI1 and MLI2 (Supplementary Figure S2A). Since Bergmann cells are one type of astrocytes, it seems reasonable that there was a high correlation between the two cell types.

Additionally, STRIDE could discover cell-type-related topics and underlying functions for these topics (Supplementary Figure S2B). Interestingly, we noticed that the topic signatures showed better spatial localization patterns compared to marker genes for molecular layers (Figure 3C, D). We also performed GO function analysis on cell-type-specific topics and demonstrated the consistency between the topics and their functions (Supplementary Figure S2C). For instance, GO terms involved in myelinogenesis, such as myelination and axon ensheathment were enriched in oligodendrocyte-specific topics (Topic 8, 18 and 30), while biological processes participating in synapse formation and synaptic signal transmission were identified in topics related to Purkinje cells and molecular layer interneurons (Topic 3 and 29). Taken conjointly, these results suggested that topic signatures generated by STRIDE could better describe the spatial localization pattern of cell types than their marker genes, which can be used to identify novel spatially expressed genes.

### STRIDE characterized the spatial heterogeneity of tumor cells in human squamous cell carcinoma microenvironment

To further explore the application of STRIDE in complex human tissues, we applied it on a hSCC dataset(15) generated using the ST technology. The scRNA-seq data from matched patients were analyzed to decipher the cell-type populations, and subsequently used as a reference to integrate with ST data. Deconvolution using STRIDE could explicitly resolve the cell-type compositions for spatial locations, thereby delineating the complex tumor microenvironment (Figure 4A). Interestingly, the deconvolution analysis by STRIDE revealed a fibrovascular niche where endothelial cells and fibroblasts were enriched, and another immune-cell-enriched leading-edge region (Figure 4A, Supplementary Figure S3A), which were in accordance with the findings from the original study. To better define spatial domains within the tissue section, we performed spatial clustering analysis combining both the neighborhood information and the cell-type deconvolution result. Locations with similar cell-type compositions and similar surrounding cell populations were clustered together (see Materials and methods for more details). This identified six spatial domains, each of which had different cell-type proportions (Figure 4B, C). Cluster C4 and C2 corresponded to the fibrovascular niche and the immune-related leading-edge mentioned above, respectively, while C3 were dominated by epithelial/malignant cells, which can be regarded as the tumor region. We next explored the relationship between the immune cell subset distribution and their relative position to tumor. According to the distance from the tumor region, we divided the spots outside C3 into two regions, the tumor-stroma border and the stroma region (Supplementary Figure S3B, see Materials and methods). We observed that there were more myeloid cells in the stroma region than in the tumor-stroma border. As for the lymphocytes, CD4^+^ T cells were enriched in the border, while B, CD8^+^ T and proliferating T cells didn’t show obvious enrichment in the border or the stroma region (Supplementary Figure S3C). This is consistent with the finding that regulatory T cells were enriched in the border region(15). Taken together, by integrating neighborhood information, STRIDE deconvolution could define the spatial domains, and further characterized the spatial distribution patterns of different cell types in the tumor microenvironment.

**Figure 4.**
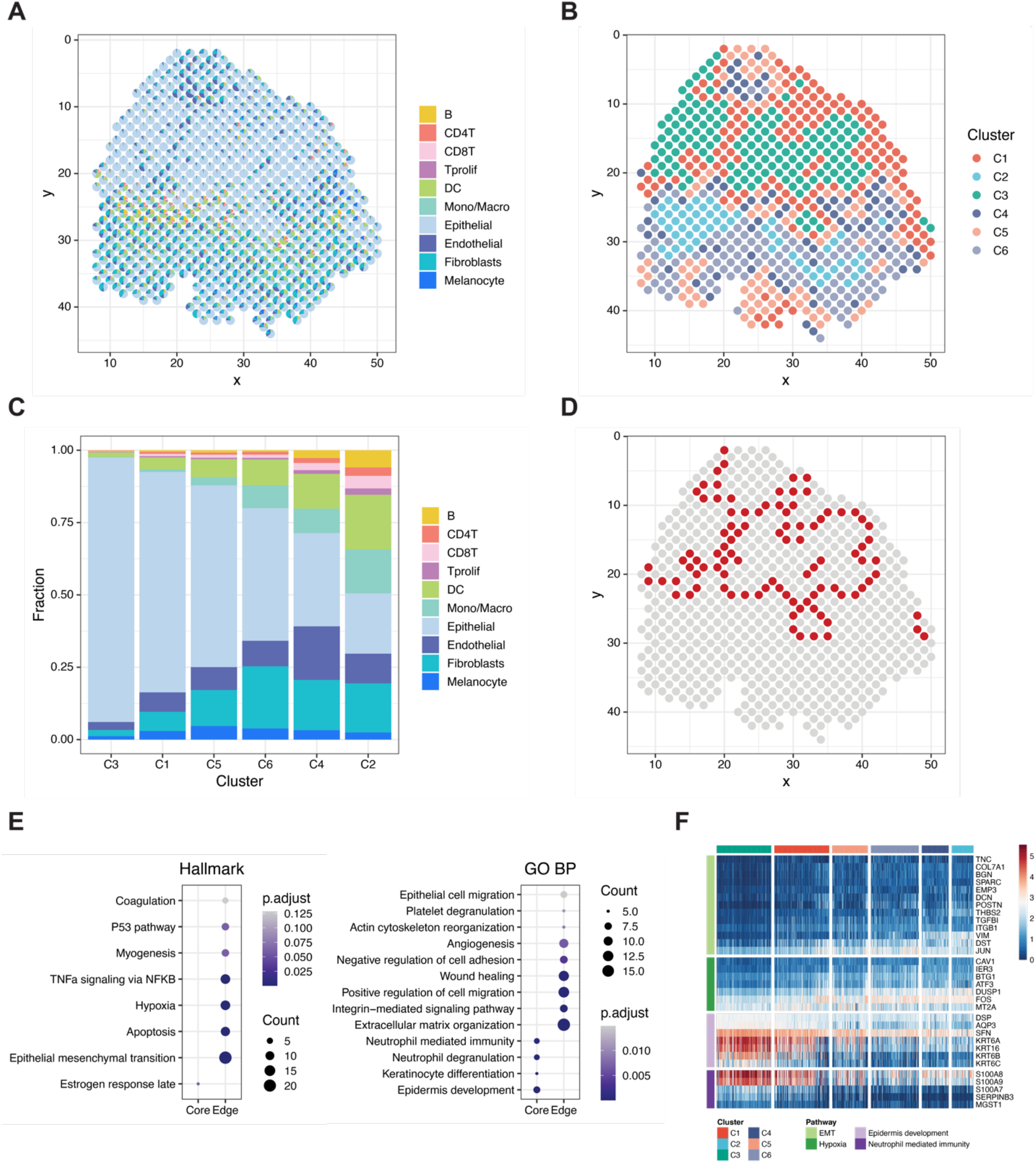
Characterizing the heterogeneity of microenvironment in human squamous cell carcinoma. A. The scatter pie plot to show the spatial locations of different cell types predicted by STRIDE. Each scatter represents a spot in the ST slide. The pie chart is used to reflect the proportions of different cell types within each spot. Colors represent different cel types. B. The k-means clustering of spots based on the cell-type compositions and surrounding cell populations. Colors represent the cluster labels. C. The cell-type composition of each cluster. The cell-type proportions of all spots in each cluster were averaged to represent the cluster’s cell-type composition. D. Spatial location of the tumor-edge region. Spots in the tumor-edge region are highlighted with red. E. Hallmark (left) and GO (right) enrichment analysis on the up-regulated genes of epithelial cells in each region. The size of the dot represents the number of genes enriched in the term, and the color represents the enrichment significance. F. The expression profile of genes associated with different pathways in epithelial cells mapped to each region. Pathways include EMT, hypoxia, epidermis development and neutrophil mediated immunity.

The tumor cells usually displayed high heterogeneity, we next asked whether the spatial domains identified by STRIDE could be used to investigate the potential relationship between tumor cells’ heterogeneity and their spatial locations. We defined C3 as tumor-core, and its surrounding region (C1, C4 and C5) as tumor-edge, respectively (Figure 4D). To describe the difference between the tumor cells from these two regions, we performed differential gene expression analysis and function enrichment analysis for the up-regulated genes of each region. Surprisingly, the tumor-core and the tumor-edge region showed distinct hallmark pathways (Supplementary Figure S3D). The tumor-core region was characterized by the enrichment of estrogen response and cholesterol homeostasis pathways, which were reported to play an important role in SCC carcinogenesis(40). By contrast, the edge region specific genes were highly enriched in interferon-involved signaling pathways, in line with the results from the previous study(15). Interestingly, we also found cancer hallmarks such as epithelial mesenchymal transition (EMT) and angiogenesis were enriched in the edge region, suggesting these tumor cells had the ability to modulate immune response and were likely to metastasize (Supplementary Figure S3D, E). To exclude the possibility that the observation was affected by other cell types present in the tumor-core and the tumor-edge region, we dissected the transcriptome of epithelial cells from the spots (see Materials and methods). Epithelial cells from scRNA-seq were mapped to the spatial locations based on the similarity of topic distribution (Supplementary Figure S3F). Again, we found that pathways involved in epidermis development and keratinization were identified in epithelial cells within the tumor-core region, while epithelial cells in the tumor-edge were enriched in pathways related to hypoxia, as well as EMT associated pathways such as cell migration and angiogenesis (Figure 4E, F). In summary, the cell-type deconvolution results from STRIDE not only helped to identify spatial localized domains, but also revealed the spatial heterogeneity within the same cell type.

### STRIDE detected the location of rare cell types during the development of human heart

To verify the utility of STRIDE in different biological systems, we also applied it to study spatial organization of organ development. A recent study presented a spatiotemporal atlas of the developing human heart by combining scRNA-seq, spatial transcriptomics and *in situ* sequencing (ISS)(14), which provided great resource for us to validate the application of STRIDE. We trained model using scRNA-seq data collected from heart at 6.5-7 post-conception weeks (PCW) (Supplementary Figure S4A), and performed cell-type deconvolution on all samples from three developmental stages (4.5-5, 6.5 and 9 PCW). As expected, the atrial and ventricular cardiomyocytes were predicted to be located in the upper and lower chambers across all three stages (Figure 5A). The epicardial cells were also correctly mapped to the outer layer of the heart, which is defined as the epicardium. Remarkably, cardiac neural crest cells and Schwann progenitor cells, a rare cell type identified by scRNA-seq, was mapped to the mesenchyme region with smooth muscle cells and fibroblast-like cells enriched (Figure 5A). This spatial localization was not achieved by the direct clustering of spots based on the gene expression profiles(14). Finally, the cell-type mapping by STRIDE was highly consistent with the spatial cell-type map (Figure 5B) created through the integration of ISS and scRNA-seq by the original study(14).

**Figure 5.**
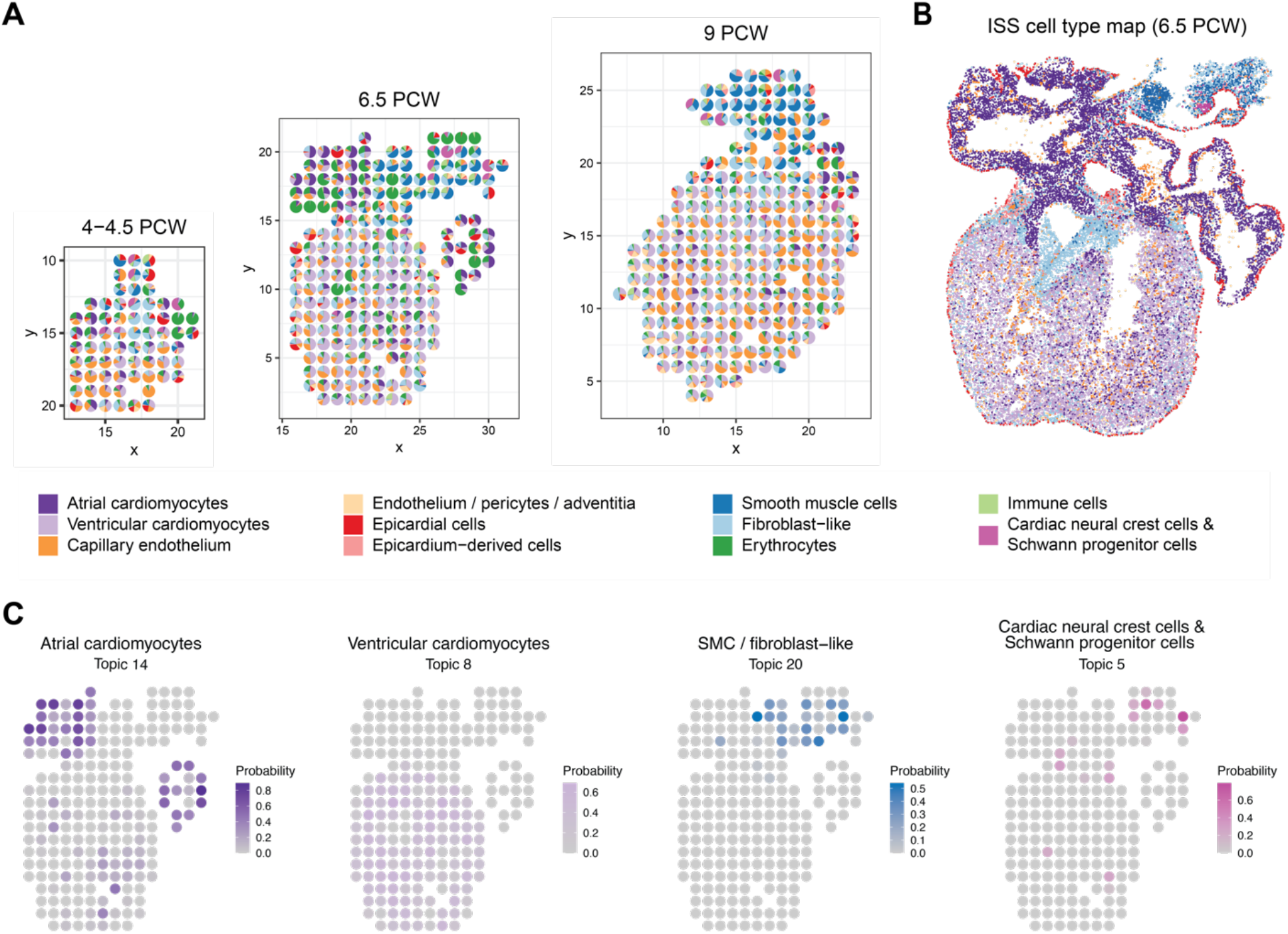
Application of STRIDE on the developing human heart. A. The deconvolution result of sample 3, 9 and 16 from 4.5-5 PCW, 6.5 PCW and 9 PCW, respectively. Each scatter represents a spot in the ST slide. The pie chart is used to reflect the proportions of different cell types within each spot. Colors represent different cell types. B. The spatial cell-type map created through the integration of ISS and scRNA-seq by the original study. C. The distribution of topics associated with atrial and ventricular cardiomyocytes, SMC or fibroblast-like cells, and cardiac neural crest cells & Schwann progenitor cells. The colors represent the probability of topics in each spot.

We further explored the distribution of topics discovered by STRIDE. Reassuringly, most cell-type-specific topics (Supplementary Figure S4B) were distributed in concordance with the spatial position of their corresponding cell types (Figure 5C). The cell-type-specific topics also showed highly cell-type-specific functions. For example, GO processes related to cardiac muscle development and contraction were enriched in cardiomyocyte-associated topics, and processes such as erythrocyte or ion homeostasis and oxidation-reduction process were enriched in erythrocyte-associated topics (Supplementary Figure S4C). Taken together, these results demonstrated that STRIDE could be used to infer the cell-type mixture patterns of tissues from different time points, and accurately detected the position even for rare cell types.

### STRIDE enabled 3D architecture construction of developing human heart using topics

In order to further demonstrate the application of STRIDE-derived topics, we set out to explore the integrative analysis across multiple samples leveraging the STRIDE deconvolution results. The human heart samples for each timepoint were sectioned along the dorsal-ventral axis, which can be integrated to reconstruct the 3D spatial architecture. We modified PASTE(35) and performed ST slide integration based on STRIDE-derived topics, which could balance both transcriptional similarity and spatial structure similarity through a balancing parameter (Figure 6A, Supplementary Figure S5A, see Materials and methods). Spots with similar topic profiles were mapped pairwise. For example, spots mainly containing ventricular cardiomyocytes were mapped to each other within the ventricle region, and the atrial cardiomyocytes in the right and left atrium were correctly mapped within a local range (Figure 6A).

**Figure 6.**
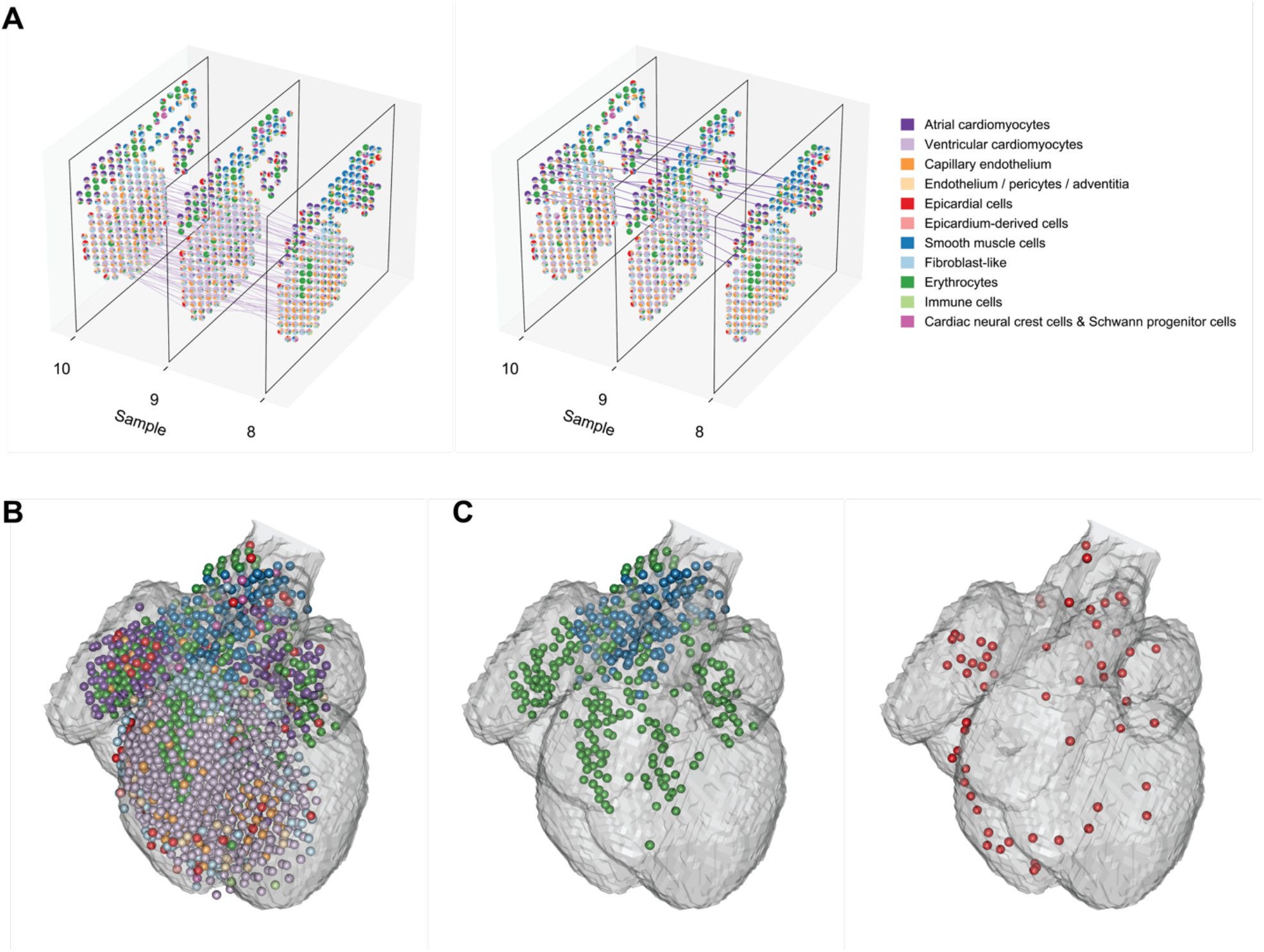
3D model reconstruction of the developing human heart. A. The alignment between adjacent tissue samples. Each pair of spots is connected by a line if they are matched according to the slide alignment. The left and right show the matching between spots dominated by ventricular and atrial cardiomyocytes, respectively. (The matching of other cell types are shown in Supplementary Figure S5A.) B. 3D model representation of the 6.5 PCW human heart constructed by STRIDE. 9 sequential samples from the 6.5 PCW heart were aligned and integrated together. Each sphere represents a spot in the ST slide, which is colored according to the cell type with the highest proportion. The translucent outline shows the 3D atlas of the developing human heart (Carnegie stage 18). C. Left, the spatial distribution of SMCs and erythrocytes. Right, the spatial distribution of epicardial cells.

We then constructed a 3D model representation of the developing human heart (Figure 6B) on the basis of sequential pairwise alignment of adjacent samples. Our 3D model could accurately represent the 3D structure only using the spatial expression information without the need for image-based registration. Interestingly, when we focused on the SMCs and erythrocytes, which are the main components of vessels, we observed that these cells started from the OFT, and clearly formed four branches into right atrium (RA), right ventricle (RV), left ventricle (LV), left atrium (LA) (Figure 6C, Supplementary Figure S5B), which might correspond to the vena cava, pulmonary artery, aorta, and pulmonary veins, respectively. In addition, we could clearly observe the location of epicardial cells in the outermost layer of the heart from the 3D model (Figure 6C). Taken together, the 3D reconstruction from multiple slides based on both the topic distribution similarity and the spatial distance similarity enabled a global view of the developing human heart, and is also applicable for other systems.

## Discussion

The recent development of spatial transcriptomics has brought new insights into the understanding of tissue architectures. However, due to the limitations of current technologies, it is difficult to accurately map cell types to spatial locations at single-cell resolution through ST data alone. To address this issue, we introduce STRIDE, a deconvolution method which utilizes topic modeling to decipher cell-type compositions of spatial transcriptomics by integrating with scRNA-seq data. Benchmarking STRIDE on different feature genes demonstrated that STRIDE could accurately estimate the cell-type proportions, and outperformed other existing methods in terms of the balance between specificity and sensitivity, and the robustness to the sequencing depth. By applying STRIDE on different scenarios, we confirmed its versatility in different biological systems and flexibility to various spatial transcriptomics techniques. We also demonstrated the broad application of STRIDE in the derivation of spatially localized cell-type signatures, identification of spatial domains, and final reconstruction of 3D tissue architecture based on cell-type-specific topics.

In spite of the aforementioned merits, STRIDE still has some limitations. First of all, the deconvolution by STRIDE depends on the assumption that the cell populations and their proportions are similar and robust between spatial transcriptomics and scRNA-seq. In fact, different sampling strategies and sites make a great difference to the cell-type compositions. It’s almost impossible to estimate the fractions of cell types which are present in spatial data but absent in scRNA-seq data. The issue can be mitigated by either controlling the consistency of sampling sites, or selecting closely matched reference samples. Secondly, STRIDE showed reduced performance in predicting the fractions of transcriptionally similar cell subtypes due to the lack of sufficient subtype-specific marker genes, although all of the other methods also failed. Future implementation of hierarchical topic modeling(41) might discover hierarchical topic structures, thus enhancing the capability to discriminate similar cell types from the same lineage. The principal objective of spatial transcriptomics is to understand the cellular heterogeneity and interplay in the spatial context. Many methods have been developed to infer the ligand-receptor interactions from spatial transcriptomics through the co-expression or co-localization analysis within the physically proximate locations(42). Future deployment of STRIDE could consider to estimate deconvolved cell-type expression level based on scRNA-seq, which could assist in the effective inference of interactions between different cell types. The coupling of cell-type distribution and intercellular interactions will facilitate the discovery of principles for the spatial organization of cells.

STRIDE could deconvolve the cell-type compositions of spatial transcriptomics based on latent topics. On the other hand, the shared topics by single-cell and spatial transcriptomics could also be used to map single cells to spatial locations. In this way, single-cell multi-omics data, such as scNOMe-seq(43), scNMT-seq(44), could be integrated with spatial transcriptomics through scRNA-seq to uncover the spatial regulatory mechanism. And with the accumulation of spatio-temporal transcriptomics and regulatory profiles, STRIDE could be further enhanced to elucidate the spatio-temporal multi-omics dynamics during the tissue development or the tumor progression. Besides, most of the current spatial techniques quantify the gene expression and infer cell-type distribution in the two-dimensional space. Though the topic-based integration of multiple spatial transcriptomics samples proved to be helpful in the 3D reconstruction of tissue architectures, we anticipate that with the aid of imaging data and data from other modalities, it will be possible to establish a more comprehensive and multi-scale 3D tissue atlas by STRIDE.

## Data availability

The mouse cerebellum dataset including spatial transcriptomics and snRNA-seq is available at https://singlecell.broadinstitute.org/single_cell/study/SCP948. The SCC dataset can be accessed from Gene Expression Omnibus (GEO) through the accession number GSE144240. The human heart dataset is collected from https://www.spatialresearch.org.

## Code availability

STRIDE is an open-source python package with source code freely available at https://github.com/wanglabtongji/STRIDE.

## Acknowledgements

This work was supported by the National Natural Science Foundation of China [32170660, 31801059, 32000561, 81872290, 81972551]. Shanghai Rising Star Program [21QA1408200]. Natural Science Foundation of Science [21ZR1467600]. The authors thank the Bioinformatics Supercomputer Center of Tongji University for offering computing resource. The authors acknowledge Junjie Hu and Yilv Yan for thr helpful discussion on the pathological annotation of H&E images.

## Author Contributions

C.W. and Q.W. conceived the project. D.S. developed the algorithm, wrote the workflow and benchmarked the performance. Z.L. helped in data processing. T.L. offered suggestions to the workflow and provided the pathological annotation of H&E images. D.S., Q.W. and C.W wrote the manuscript. C.W supervised the whole project. All authors read and approved the final manuscript.

