## Supplementary Figures for "STRIDE: accurately decomposing and integrating spatial transcriptomics using single-cell RNA sequencing"

### Supplementary Figure S1

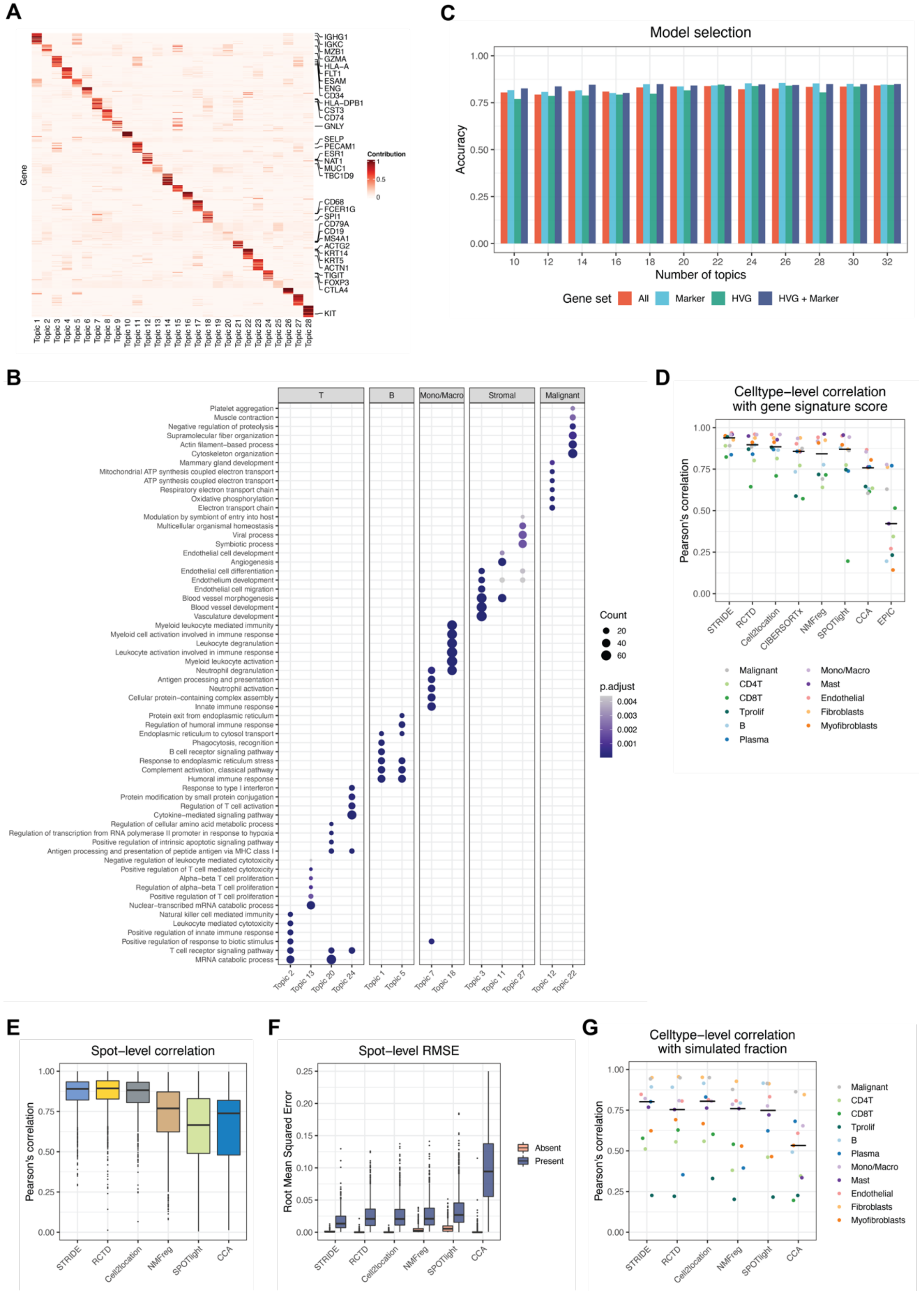

**Benchmarking STRIDE's performance using simulated data**

A. The gene-by-topic distribution estimated by STRIDE. For each topic, the top 20 genes with the highest contributions are shown, and the cell-type-specific marker genes are marked on the right.

B. GO function analysis on the topics derived from STRIDE. For each topic, GO analysis was performed on the top 200 genes with the highest contributions. The topics are grouped by their associated cell types. The size of the dot represents the number of genes enriched in the term, and the color represents the enrichment significance.

C. The accuracy of cell-type assignment for single cells for different topic numbers. For each gene set, the optimal topic number was selected by the highest accuracy.

D. Benchmarking the ability to distinguish diverse cell types across different deconvolution methods. Pearson's correlation between the predicted proportions and gene signature score was calculated for each cell type. The black line in each column indicates the median of different cell-types' correlation for each method.

E. Benchmark of STRIDE's accuracy against different deconvolution methods on the simulated dataset with correlated cell-type distribution. The box plot reflects the overall distribution of Pearson's correlation calculated in each spot for each method.

F. Benchmark of STRIDE's sensitivity and specificity against different deconvolution methods. All the simulated locations were divided into two groups according to the presence (blue) and absence (pink) of each cell-type, and RMSE was calculated within each group. The box plot reflects the distribution of RMSE in different methods.

G. Benchmark of the ability to distinguish diverse cell types across different deconvolution methods. Pearson's correlation between the predicted proportions and the ground truth was calculated for each cell type. The black line in each column indicates the median of different cell-types' correlation for each method.

**Supplementary Figure S2**

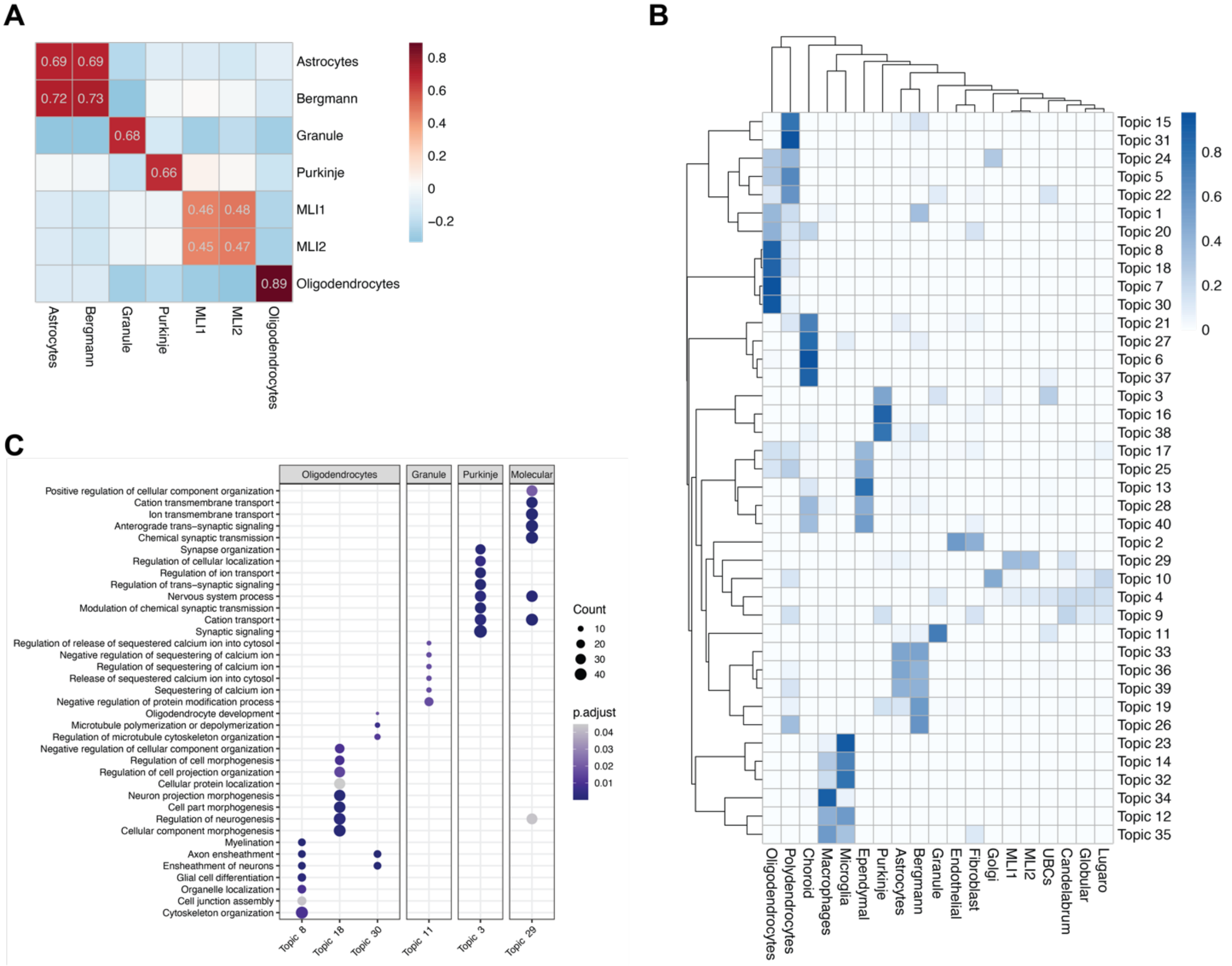


**Application of STRIDE on the mouse cerebellum**

A. Pearson's correlation between the predicted proportions and gene signature score across different cell types.

B. The cell-type-by-topic distribution estimated by STRIDE. The color represents the probability that one topic exists in one given cell type.

C. GO function analysis on the topics derived from STRIDE. For each topic, GO analysis was performed on the top 200 genes with the highest contribution. The topics are grouped by their associated cell types. The size of the dot represents the number of genes enriched in the term, and the color represents the enrichment significance.

#### Supplementary Figure S3

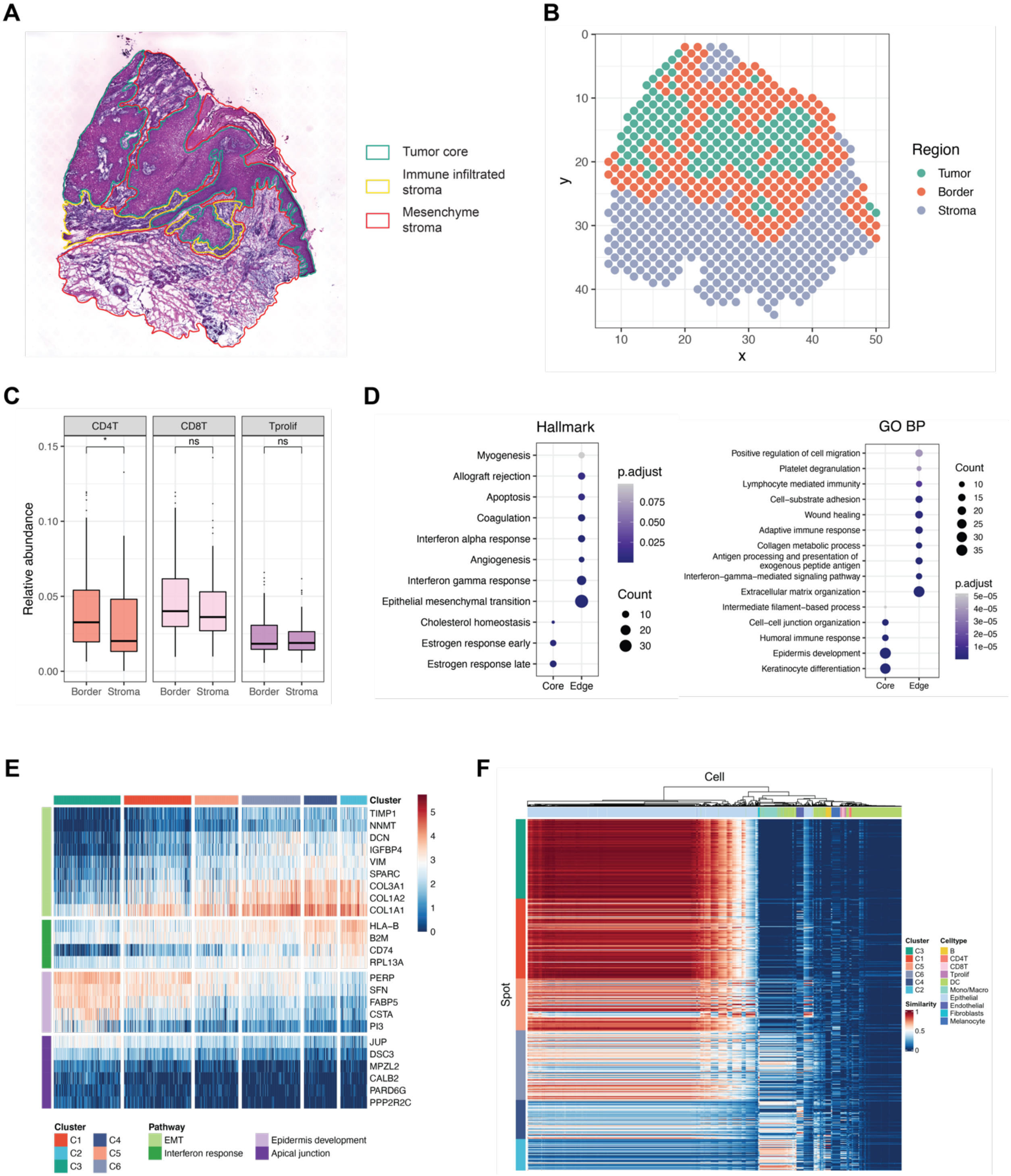

**Characterizing the heterogeneity of microenvironment in human squamous cell carcinoma**

A. The H&E staining of the hSCC sample. Full lines with different colors are used to label the pathological annotation results.

B. Spatial location of the tumor, tumor-stroma border and stroma region.

C. The difference of the distribution of immune cells within the border and stroma region. The y axis represents the relative abundance of cell types. ‘ns’ represents p-value > 0.05, ‘*’ represents 0.01 < p-value <= 0.05, and ‘****’ represents p-value <= 0.0001.

D. Hallmark (left) and GO BP (right) enrichment analysis on the up-regulated genes of spots in each region. The size of the dot represents the number of genes enriched in the term, and the color represents the enrichment significance.

E. The expression profile of genes associated with different pathways in spots within each region. Pathways include EMT, interferon response, epidermis development and apical junction.

F. The topic-based cosine similarity between spots and cells. The rows and the columns indicate spots from spatial transcriptomics and cells from scRNA-seq, respectively.

#### Supplementary Figure S4

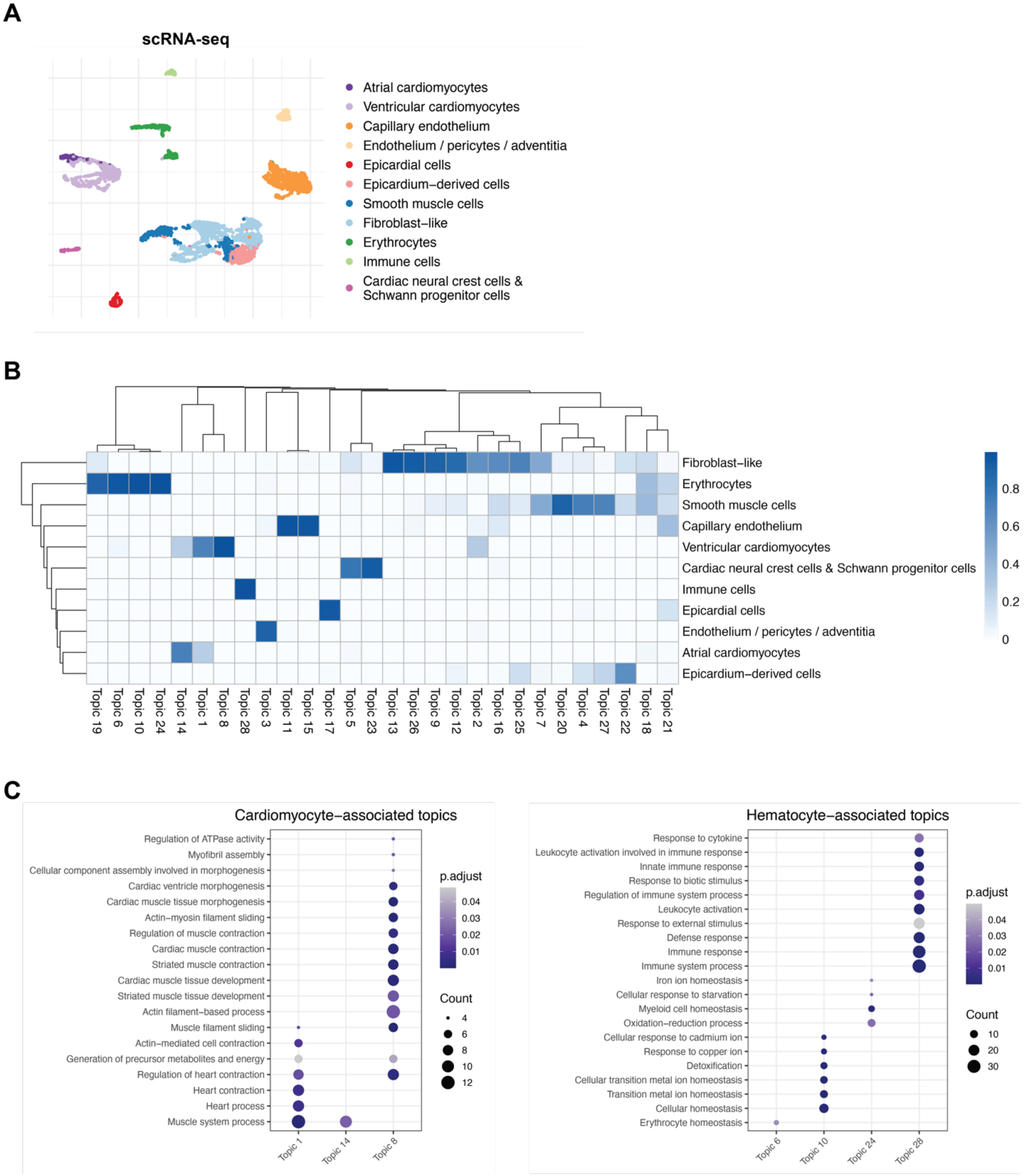

**Application of STRIDE on the developing human heart**

A. UMAP plot of paired scRNA-seq. Cells are colored according to the cell-type annotations from the original paper.

B. The cell-type-by-topic distribution estimated by STRIDE. The color represents the probability that one topic exists in one given cell type.

C. GO function analysis on the topics associated with cardiomyocytes (left) and hemocytes (right). For each topic, GO analysis was performed on the top 50 genes with the highest contribution. The size of the dot represents the number of genes enriched in the term, and the color represents the enrichment significance.

#### Supplementary Figure S5

**A**

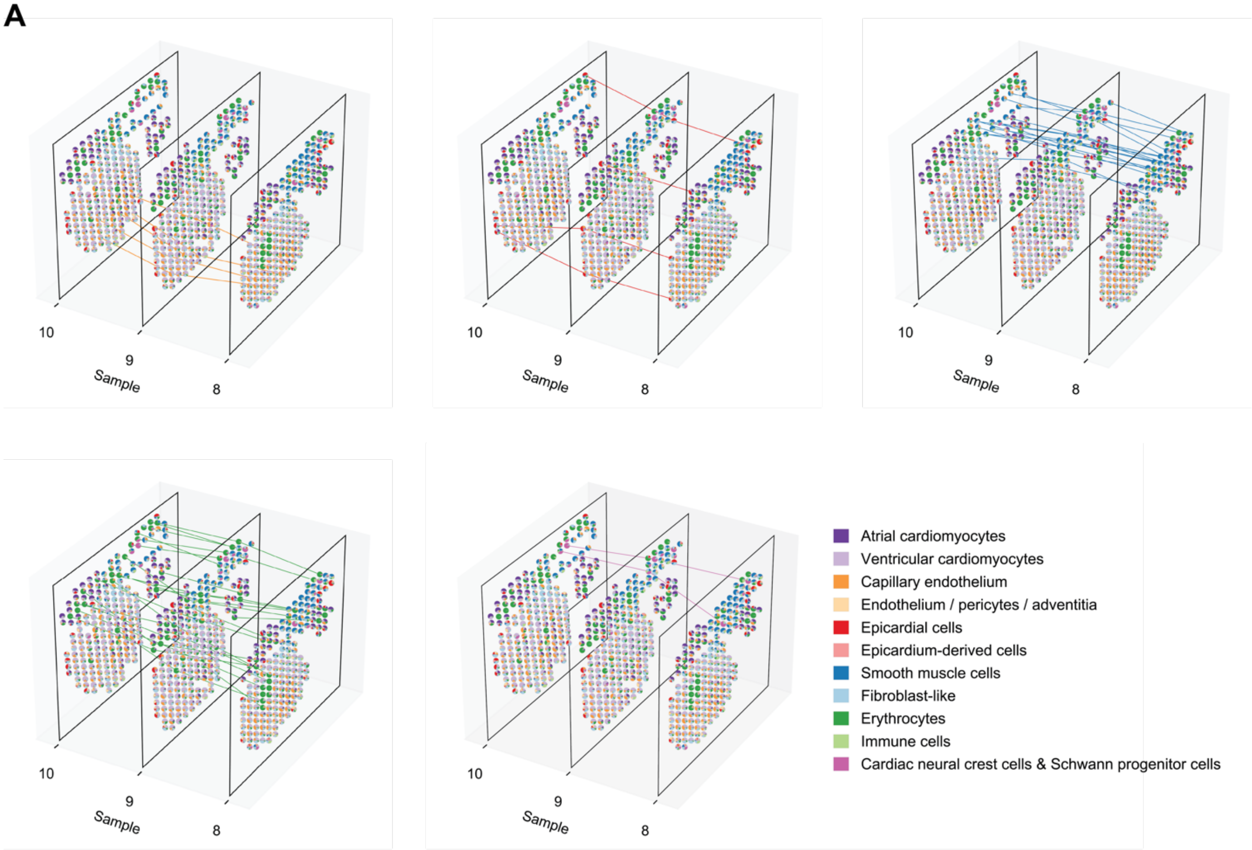


**B**

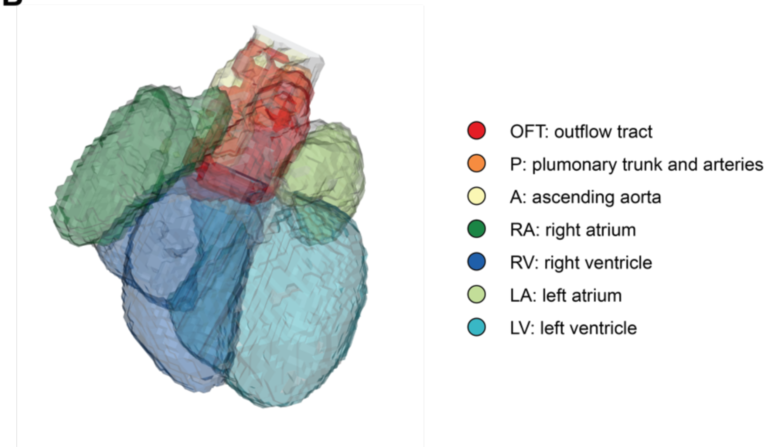


**3D model reconstruction of the developing human heart**

A. The alignment between adjacent tissue samples. Each pair of spots is connected by a line if they are matched according to the slide alignment.

B. 3D atlas of the developing human heart (Carnegie stage 18). Different anatomical regions are painted with different colors. OFT: outflow tract; P: pulmonary trunk and arteries; A: ascending aorta; RA, right atrium; RV, right ventricle; LA, left atrium; LV, left ventricle.
